# Transcriptomic analysis of probable asymptomatic and symptomatic Alzheimer brains

**DOI:** 10.1101/621912

**Authors:** Hamel Patel, Angela K. Hodges, Charles Curtis, Sang Hyuck Lee, Claire Troakes, Richard J.B Dobson, Stephen J Newhouse

## Abstract

Individuals with intact cognition and neuropathology consistent with Alzheimer’s disease (AD) are referred to as asymptomatic AD (AsymAD). These individuals are highly likely to develop AD, yet transcriptomic changes in the brain which might reveal mechanisms for their AD vulnerability are currently unknown. Entorhinal cortex, frontal cortex, temporal cortex and cerebellum tissue from 27 control, 33 AsymAD and 52 AD human brains were microarray expression profiled. Differential expression analysis identified a significant increase of transcriptomic activity in the frontal cortex of AsymAD subjects, suggesting fundamental changes in AD may initially begin within the frontal cortex region prior to AD diagnosis. Co-expression analysis identified an overactivation of the brain “glutamate-glutamine cycle”, and disturbances in the brain energy pathways in both AsymAD and AD subjects, while connectivity of key hub genes in this network indicates a shift from an already increased cell proliferation in AsymAD subjects to stress response and removal of amyloidogenic proteins in AD subjects. This study provides new insight into the earliest biological changes occurring in the brain prior to the manifestation of clinical AD symptoms and provides new potential therapeutic targets for early disease intervention.

## Introduction

The increase in life expectancy has profoundly increased the ageing population, which, unfortunately, is also accompanied by a rise in age-related disorders including Alzheimer’s disease (AD) [1]. Alzheimer’s disease is a neurodegenerative disorder characterised by progressive accumulation of extracellular amyloid-β (Aβ) protein and intracellular hyperphosphorylated tau filaments in the brain, which form insoluble plaques and tangles respectively. These protein aggregates affect neuronal activity which can lead to progressive loss of neurons associated with deterioration in cognition and development of neuropsychiatric symptoms.

Through longitudinal studies involving autopsy, it has become evident that clinical signs of cognitive impairment are apparent after substantial years of neurodegeneration, which occurs decades after neuropathological changes [2]. As the disease is progressively slow and as everyone is expected to experience cognitive change during normal ageing, differentiating AD symptoms from normal ageing at an early stage of disease can be difficult. Up to 20-30% of the ageing population with intact cognition have amyloid deposition, with these individuals at higher risk of progressing to AD than those without amyloid [3]. These individuals are often referred to as asymptomatic AD (AsymAD) [4] and have been shown to be distinguishable from normal ageing based on neuropathology, brain imaging and cerebrospinal fluid biomarkers [2]. While some of these individuals progress to developing symptoms related to cognition, which deviate from normal Mild Cognitive Impairment (MCI), and then to AD, not all do. They are therefore a heterogeneous group, representing those with prodromal AD and those impervious to AD despite having the pathological hallmarks.

Measuring genome-wide expression of transcripts as markers of gene activity has revealed that cognitive decline is accompanied by changes in brain gene expression from normal ageing through to MCI and AD. Studies have suggested that some changes in the pattern of gene expression in normal ageing such as synaptic function and energy metabolism [5], are extensively altered in MCI [6] and AD [7] [8] [9] [10] [11]. Additional work has also suggested a number of other biological pathways are more specifically altered in AD, including inflammation [10] [12] [11], protein misfolding [10] [12], transcription factors [10] [12], cell proliferation [10] [12], immune response [13] [14] [15] [16], protein transcription/translation regulation [13] [14] [17] [18] [19] [20], calcium signalling [13] [21] [9], MAPK signalling [19] [8], and various metabolism pathways [19] [22] [23] [24] [20] [14] [25] which reflect the extent and type of pathology and disruption to cell activity as disease progresses. It is unknown how early the different types of changes occur in the brain, such as in the pre-symptomatic phase or specifically in AsymAD subjects who already have the pathological hallmarks of AD such as amyloid and neurofibrillary tangles (NFTs). Understanding the fundamental changes in this AsymAD group may shed light on specific biological mechanisms that may be involved in early pathological hallmarks of AD, providing new therapeutic targets for early intervention.

In this study, we investigated transcriptomic changes in the human brain of healthy ageing, AsymAD and AD subjects, which have been classified based on clinical assessment before death and AD neuropathology at autopsy. Typical transcriptomic analysis coupled with a systems-biology approach was used to identify disturbances in the underlying biological mechanisms across the entorhinal cortex, temporal cortex, frontal cortex and cerebellum brain regions. In addition, we provide access of gene-level results to the broader research community through a publicly available R SHINY web-application (https://phidatalab-shiny.rosalind.kcl.ac.uk/ADbrainDE), allowing researchers to quickly query the expression of specific genes through the progression of AD and across multiple brain regions.

## Materials and Methods

### Medical Research Council London Neurodegenerative Diseases Brain Bank

A total of 112 brains were obtained from the Medical Research Council (MRC) London Neurodegenerative Diseases Brain Bank (from now on referred to as MRC-LBB) hosted at the Institute of Psychiatry, Psychology and Neuroscience, KCL. All cases were collected under informed consent, and the bank operates under a licence from the Human Tissue Authority, and ethical approval as a research tissue bank (08/MRE09/38+5). Neuropathological evaluation for neurodegenerative diseases was performed in accordance with standard criteria.

### MRC-LBB sample selection

BRAAK staging is a measure of the spread of hallmark AD pathology across the brain and is part of the neuropathological assessment. In general, BRAAK stages I-II, III-IV and V-VI have been suggested to represent prodromal, early-moderate AD, and moderate-late AD respectively. Twenty-seven control cases were used - classified as showing no clinical sign of any form of dementia and no neuropathological evidence of neurodegeneration. Thirty-three AsymAD cases were also analysed - defined as clinically dementia-free at the time of death, but neuropathological assessment at autopsy showed hallmark AD pathology. Finally, fifty-two AD cases, which had both a clinical diagnosis of AD at death and confirmation of this diagnosis through neuropathological evaluation at autopsy, were selected.

### MRC-LBB brain region selection and RNA extraction

Frozen tissues (0.5-1cm3) from the following brain regions from each case were macrodissected into RNAlater RNA Stabilization Reagent (Qiagen): 1) Frontal Cortex (FC), 2) Temporal Cortex (TC), 3) Entorhinal Cortex (EC) and 4) Cerebellum (CB). Hallmark AD pathology was confirmed in the entorhinal cortex, temporal cortex and frontal cortex but absent from the cerebellum of AsymAD and AD subjects. RNA extraction was performed within 24 hours of dissection. Total RNA was extracted using RNeasy Lipid Tissue Mini Kit (Qiagen,74804) following the manufacturer’s protocol. Genomic DNA was removed using gDNA Eliminator Spin Columns (Qiagen). RNA quality was evaluated with an Agilent 2100 bioanalyzer (Agilent Technologies, Inc., Palo Alto, CA).

### MRC-LBB Illumina beadArray expression profiling

Total RNA (25ng) was prepared for array expression profiling using the Ovation Pico WTA system (NuGEN Technologies, Inc., San Carlos, CA), as described by the manufacturer’s protocol. The Nugen system is optimised for the amplification of degraded RNA, where amplification is initiated at the 3’ end as well as randomly throughout the whole transcriptome. The samples were processed at the NIHR Biomedical Research Centre for Mental Health (BRC-MH), Genomics & Biomarker Core Facility at the Social, Genetic & Developmental Psychiatry Centre, Institute of Psychiatry, Psychology and Neuroscience, King’s College London (https://www.kcl.ac.uk/ioppn/depts/sgdp-centre/research/The-IoPPN-Genomics--Biomarker-Core-Facility.aspx) in accordance with the manufacturer’s protocol using the Illumina HT-12_V4 beadchips (Illumina, USA).

### Microarray expression data processing

Raw gene expression data was exported from Illumina’s GenomeStudio (version 2011.1) into RStudio (version 0.99.467) for data processing. Using R (version 3.2.2), raw data was Maximum Likelihood Estimation (MLE) background corrected using R package “MBCB” (version 1.18.0) [26], log2 transformed, and underwent Robust Spline Normalisation (RSN) using R package “lumi” (version 2.16.0) [27].

A series of quality control steps were carried out before data analysis. Duplicate samples were removed based on lowest RIN score. Sex was predicted for each sample using the R package “massiR” (version 1.0.1) [28], with any discrepancies in predicted and clinically recorded sex from the same individual across all tissues removed from further analysis. For each sample, probesets “not reliably detected” or “unexpressed” were removed to eliminate noise [29] and increase power [30]. If the expression of a probe was below the 90^th^ percentile of the log2 expression scale in over 80% of samples across all groups (based on disease status, brain region and sex), the probe was deemed “unexpressed” and was removed from further analysis.

Batch effects were then explored using Principal Component Analysis (PCA) and Surrogate Variable Analysis (SVA) using the R package “sva” (version 3.10.0) [31]. Sex and diagnosis information was used as covariates in sva when correcting for unknown batch effects. To ensure homogeneity among the biological groups, outlying samples per tissue and disease group were iteratively identified and removed following the fundamental network concepts described in [32]. Finally, Illumina-specific probe ID’s were converted to the universal Entrez Gene ID using the R package “illuminaHumanv4.db” (version 1.22.1).

### Differential Expression and Gene Set Enrichment Analysis

Differential Expression (DE) analysis was performed using the R package “limma” (version 3.20.9) [33]. As we had theoretically corrected for unwanted batch effects in our data using sva, we only used sex in the DE model as a covariate. A gene was regarded as significantly differentially expressed if the false discovery rate (FDR) adjusted p-value ≤ 0.05.

Gene set enrichment analysis (GSEA) was performed using an Over-Representation Analysis (ORA) implemented through the ConsensusPathDB (http://cpdb.molgen.mpg.de) web-based platform (version 32) [34] in October 2017. ConsensusPathDB incorporates numerous well-known biological pathway databases including BioCarta, KEGG, Reactome and Wikipathways. It performs a hypergeometric test while combining a background gene list, compiles results from each database and corrects for multiple testing using FDR [34]. During GSEA analysis, a minimum overlap of the query signature and database was set as 2.

### Weighted Gene Co-expression Network Analysis

Weighted gene co-expression analysis (WGCNA) was performed using R package “WGCNA” (version 1.51) to identify clusters (modules) of highly correlated genes, with the underlying hypothesis that such modules could possess a common function. The WGCNA analysis was performed as described in [35]. In brief, a co-expression network based on “signed” adjacency was independently created for all three phenotypes (control, AsymAD and AD group), topological overlap calculated, and hierarchical clustering used to group genes into modules. The control group module was assigned default colours based on module size, and the AsymAD and AD module colours determined based on the control module gene overlap. Module cross-tabulations were generated across the three phenotypes and Fisher’s exact test used to test for enrichment between modules-gene assignments between the control, AsymAD and AD groups. To aid in identifying significant changes in the co-expression network within the same modules in the three phenotypes, additional statistics known as “Module preservation Zsummary” and “median rank” were calculated as described in [36].

### Protein-Protein Interaction network analysis

Protein-protein interaction (PPI) networks were generated by uploading gene lists (referred to as seeds in network analysis) to NetworkAnalyst’s (http://www.networkanalyst.ca/faces/home.xhtml) web-based platform in December 2017. The “zero-order network” option was incorporated to allow only seed proteins directly interacting with each other, preventing the well-known “hairball effect” and allowing for better visualisation and interpretation [37]. Sub-modules with a p-value ≤ 0.05 based on the “InfoMap” algorithm [38] were deemed significant “hubs” and the gene(s) with the most connections within this network as the “key hub gene(s)”.

### Study design

Differential and co-expression analysis was performed between the three disease groups and for each of the four brain regions. First, the control and AsymAD groups were compared, and from this point onwards is referred to as the “**Early AD**” analysis. Second, the AsymAD and AD groups were compared, and from this point onwards is referred to as the “**Late AD**” analysis. Finally, the control and AD groups were compared, and from this point onwards is referred to as the “**Standard AD**” analysis. An overview of the study design and analyses is shown in Figure 1.

**Figure 1:**
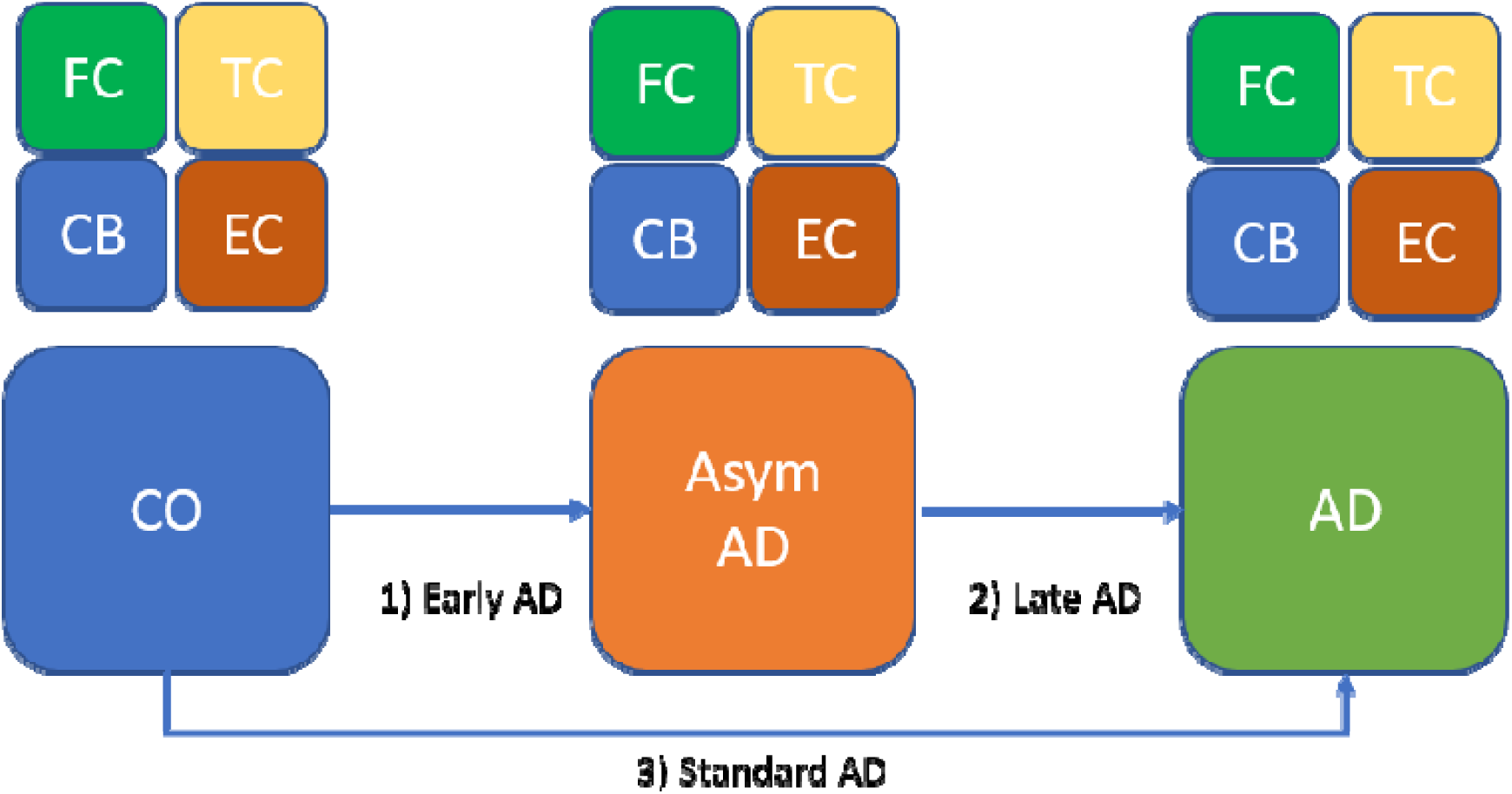
Overview of Study Design. Four brain regions; frontal cortex (FC), temporal cortex (TC), entorhinal cortex (EC) and cerebellum (CB) from the three subject groups; control (CO), Asymptomatic AD (AsymAD) and Alzheimer’s disease (AD) were expression profiled. The typical comparison between the CO and AD group is referred to as the “Standard AD” analysis, the comparison between the CO and AsymAD group is referred to as the “Early AD” analysis and the comparison between the AsymAD and AD group is referred to as the “Late AD” analysis.

### Data availability

The microarray data has been deposited in NCBI’s GEO database under the accession number GSE118553. Additionally, a shiny application was written in R using the “shiny” framework (version 0.14) to allow quick visualisation of specific gene expression in the control, AsymAD and AD subjects, and across the EC, TC, FC and CB brain regions. The application also displays DE results of each gene and can be accessed at https://phidatalab-shiny.rosalind.kcl.ac.uk/ADbrainDE. All data analysis scripts used in this study are available at https://doi.org/10.5281/zenodo.1400644

## Results

### Data processing

Of the 401 tissue samples assessed (extracted from the 112 brains) 48 samples were removed due to duplication, 4 samples due to outlier detection analysis and 2 samples due to sex discrepancies between recorded and actual sex, leaving 347 tissue samples from 111 brains for DE and co-expression analysis. As a result of samples not being microarray profiled due to sample quality, and samples being removed during the Quality Control (QC) process, not all subjects had tissue samples extracted from all four brain regions. The demographics for datasets by brain region and sample group is provided in Table 1.

**Table 1:**
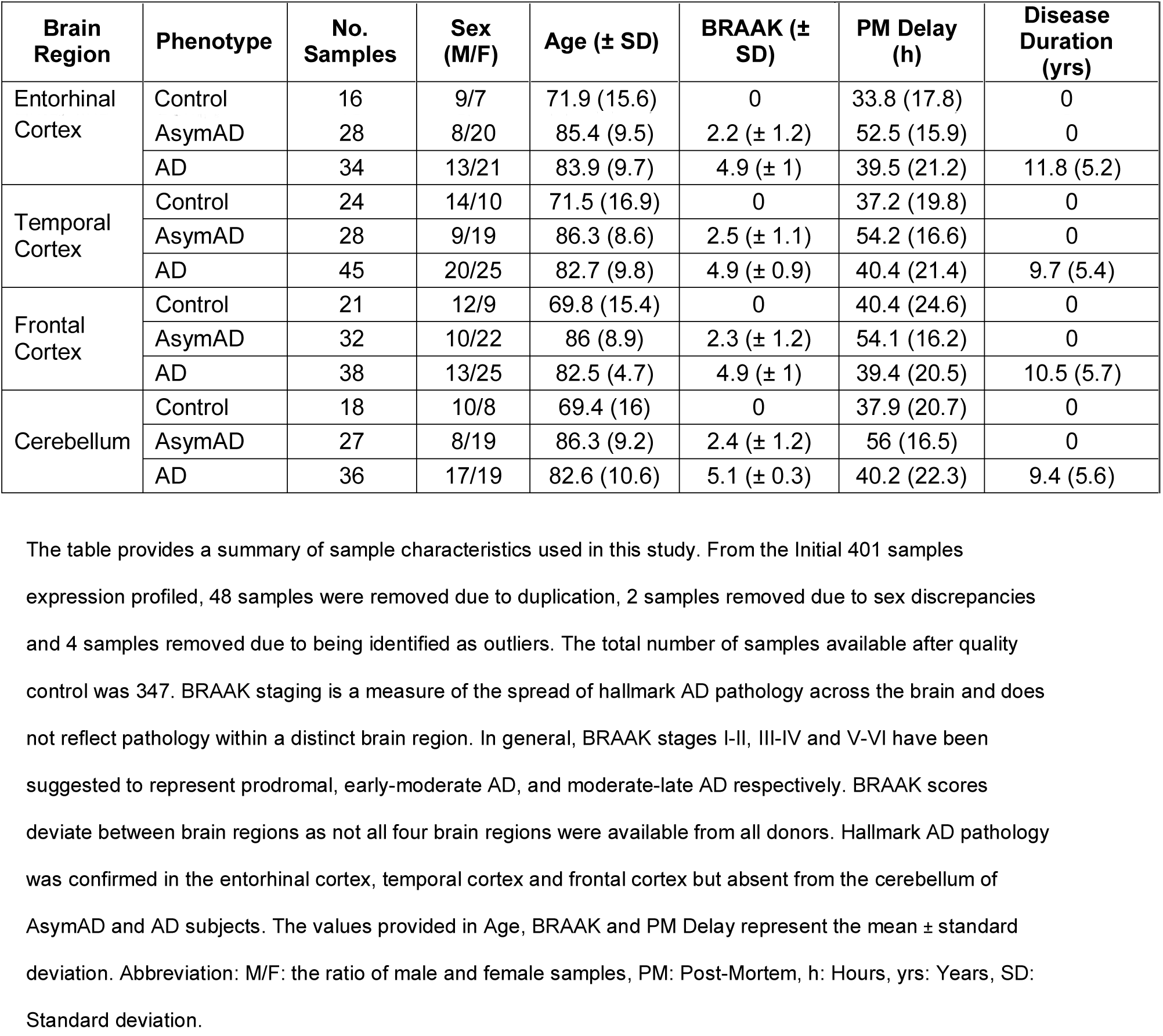
Summary of MRC-LBB sample characteristics

After further QC and annotation to determine Entrez gene identifiers, the final data represented 3518 “reliably detected” genes across all samples. Chi-squared tests revealed no significant difference in the proportion of males to females across the three disease groups or brain regions. Mann-Whitney U test revealed no significant difference between post-mortem (PM) delay or disease duration across analyses, however, age was significantly (p ≤ 0.01) lower in the control groups when compared to the AsymAD and AD group in each tissue (see Supplementary Table 1). Detailed phenotype per sample is provided in Supplementary Table 2.

### Summary of differentially expressed genes across disease groups and tissues

A summary of DEG’s identified in each brain tissue and analyses are illustrated in Figure 2, and a full list of DEG’s is provided in Supplementary Table 3. The general trend of DEG’s in subjects with AD (“Late AD” and “Standard AD” analysis) decreases across brain regions in the order of EC (n=1904 and n=1690 respectively) > TC (n=1546 and n=1517 respectively) > FC (n=52 and n=299 respectively) > CB (n=13 and n=176 respectively). This expression pattern corresponds to the route AD pathology is seen to spread through the brain. By contrast, the pattern differs in the AsymAD group (“Early AD” analysis), where most DEGs are detected in the FC (n=398) followed by the TC (n=253), EC (n=19) and CB (n=1), suggesting initial molecular changes may begin in the FC brain region prior to AD symptoms.

**Figure 2:**
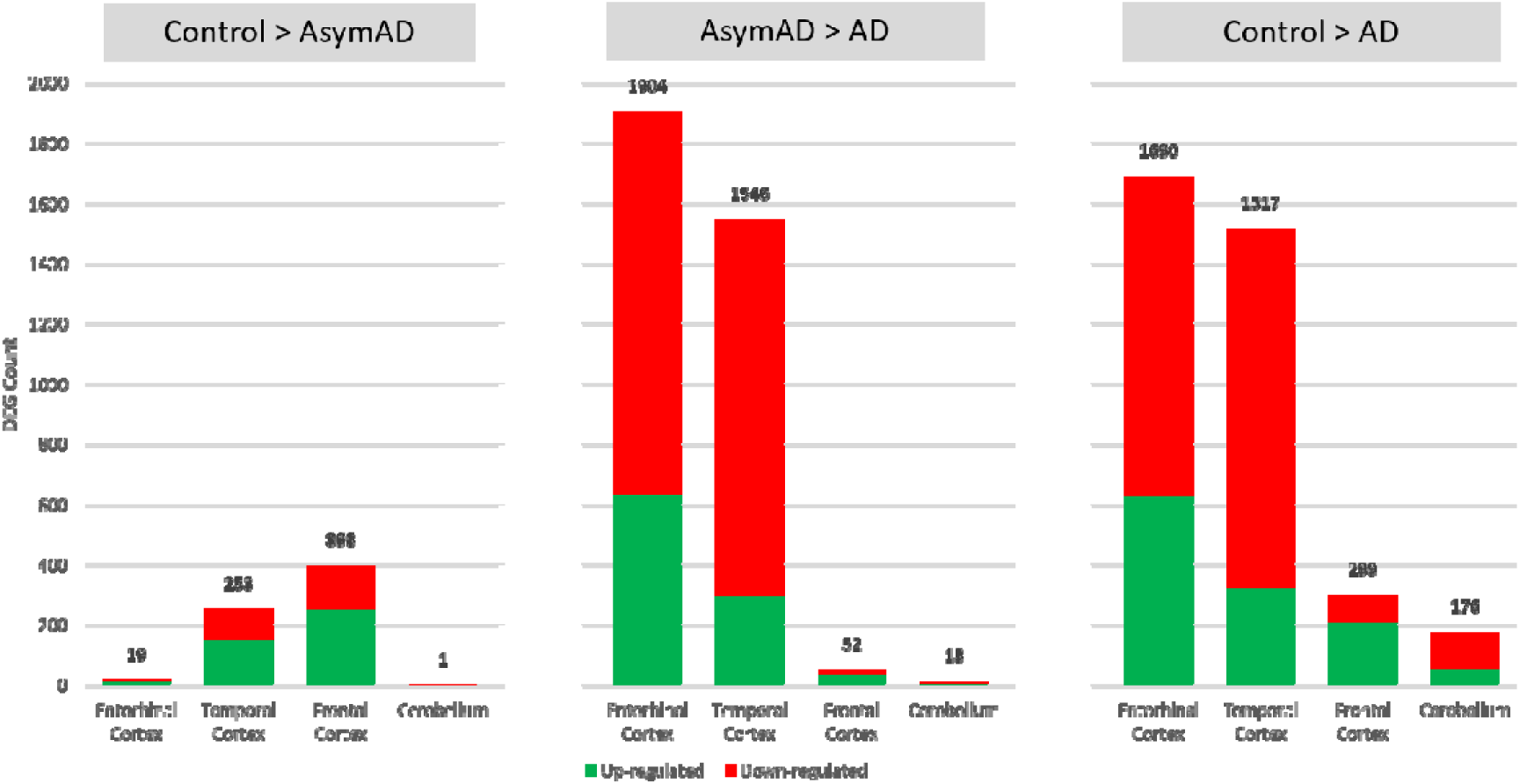
Distribution of Significant DEG (FDR adjusted p-value ≤ 0.05) across brain regions and analyses. “Control>AsymAD” summarises the number of DEGs between the control and AsymAD group. “AsymAD>AD” summarises the number of DEGs between the AsymAD and AD group. “Control>AD” summarises the number of DEGs between the control and AD group. The proportion of up-regulated genes is represented in green while the down-regulated genes are represented in red. The total number of significantly differentially expressed genes in each brain region and analysis is provided on top of each bar. More genes are observed to be generally perturbed when comparing the AD group to the AsymAD or healthy ageing group, with the general pattern of more genes perturbed in the entorhinal cortex, followed by the temporal cortex, frontal cortex and then the cerebellum, a pattern generally representing the spread of hallmark AD pathology. In contrast, comparing the AsymAD group to the healthy ageing group reveals more genes are perturbed in the frontal cortex, followed by the temporal cortex, entorhinal cortex and then the cerebellum, suggesting initial molecular changes in AD may begin in the frontal cortex before the manifestation of clinical AD symptoms.

### AD tau pathology marker suggests AsymAD subjects are an Intermediate state between normal ageing and AD

A previous study identified eight genes highly correlated with AD tau pathology [39], of which two genes (**RELN, TRIL**) are present in our data. DE analysis results indicate the **TRIL** gene expression gradually increases through the control, AsymAD and then the AD group. In addition, the expression increase is only observed in brain regions known to be affected by tau pathology (EC, TC and FC), and the extent of expression change within these affected brain regions shadows the route of disease manifestation through the brain (Figure 3a). The EC exhibits the most significant increase of **TRIL** expression (logFC=0.99, FDR adjusted p-value=2.77e-8), followed by the TC (logFC=0.48, FDR adjusted p-value=1.41e-3) and then FC brain region (logFC=0.44, FDR adjusted p-value=2.21e-2). This expression pattern further suggests the **TRIL** gene is a reliable brain marker for tau pathology, and our AsymAD samples are a good representation of early-intermediate state between normal ageing and AD.

**Figure 3:**
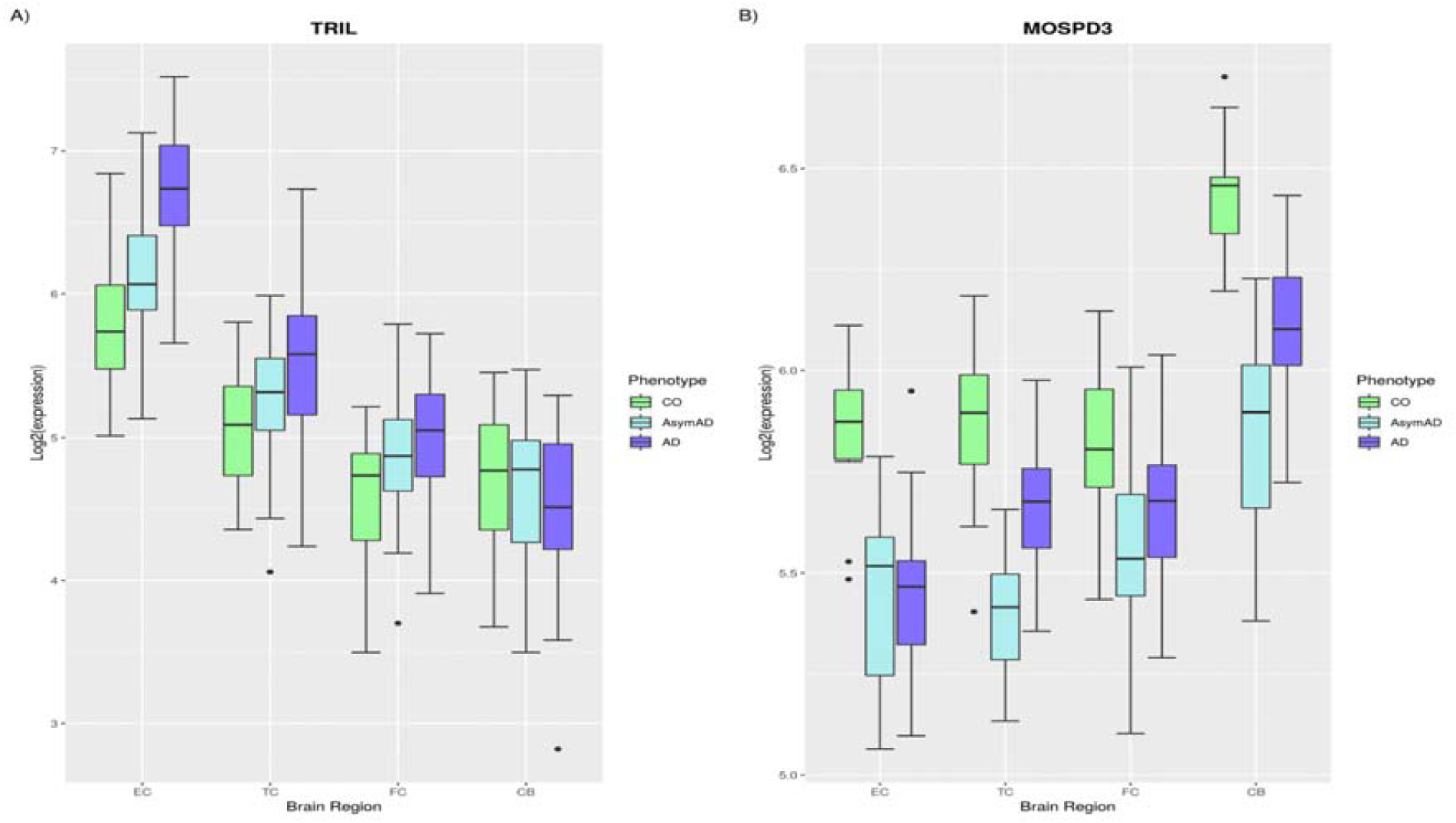
Expression boxplots of the TRIL and MOSPD3 genes. A) Previous research identified the TRIL gene as a marker for Tau pathology. The TRIL gene is significantly up-regulated (before multiple correction) from control to AsymAD (EC: logFC= 0.39 & p-value=0.01, TC: logFC=0.24 & p-value=0.04, FC: logFC=0.24 & p-value=0.04) and then further to AD (EC: logFC= 0.6 & p-value=6.57e-6, TC: logFC=0.29 & p-value=4.19e-3, FC: logFC=0.18 & p-value=0.05), but not in the cerebellum (control to AsymAD: logFC= −0.01 & p-value=1, AsymAD to AD: logFC=−0.19 & p-value=0.2), a region spared by hallmark AD pathology. The expression pattern of the TRIL gene further supports the assignment of AsymAD samples, which were based on clinical records and neuropathological assessment, as an early intermediate state between healthy ageing and AD. B) The MOSPD3 gene is the most Significant DE gene in the Early AD analysis and is consistently down-regulated in all brain regions of the AsymAD group when compared to controls (EC: logFC=−0.38 & adjusted p-value=5.6e-4, TC: logFC=−0.5 & adjusted p-value=1.18e-10, FC: logFC= −0.27 & adjusted p-value=6.91e-4, CB: logFC=−0.59 & adjusted p-value= 1.51e-6). As all brain regions are affected in AD, albeit not to the same degree, the MOSPD3 gene may represent an early brain biomarker for cell dysfunction in AD.

### The most significant differentially expressed genes per analysis

The most DEG’s from each analysis is 1) **MOSPD3** (downregulated in the TC brain region in “Early AD”, FDR adjusted p-value = 1.18e-10, Figure 3b), 2) **NPC2** (upregulated in the EC brain region in the “Late AD” analysis, FDR adjusted p-value = 2.39e-20, available to view in the SHINY web-app) and 3) **NOTCH2NL** (upregulated in the EC brain region in the “Standard AD” analysis, FDR adjusted p-value = 1.29e-15, available to view in the SHINY web-app).

### Common differentially expressed genes across all brain regions

The overlap of DE genes across brain regions is shown in Figure 4. **MOSPD3** is the only gene significantly differentially expressed across all four brain regions in the “Early AD” analysis. No gene was significantly differentially expressed in the “Late AD” analysis across all four brain regions; however, six genes (**NPC2, DUSP1, GPM6B, SLC38A2, ANKEF1, MOSPD3**) were identified in “Standard AD” analysis. Three of these genes (**DUSP1, SLC38A2** and **MOSPD3**) are consistently expressed in the same direction across all four brain regions. **DUSP1** and **SLC38A2** gene expression are upregulated during disease progression (Control to AsymAD to AD). **MOSPD3**, however, is downregulated in the disease in both the “Early AD” and “Standard AD” analyses, with no significant difference between the AsymAD and AD subjects. The remaining three genes (**NPC2, GPM6B, ANKEF1**) are DE in the same direction across all brain regions but reversed in the CB; a brain region suggested to be spared by hallmark AD pathology.

**Figure 4:**
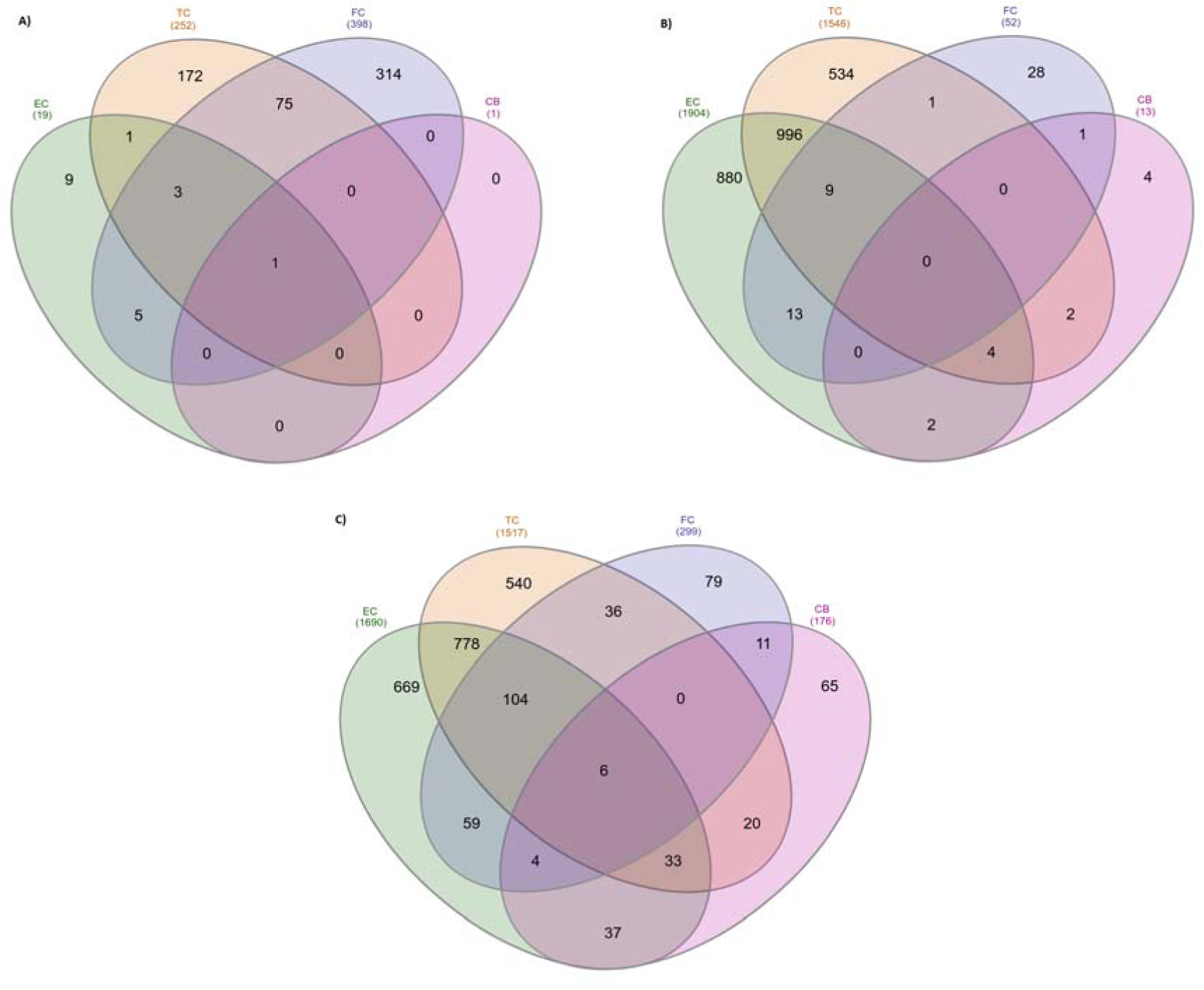
Overlap of significant DEG across brain regions in A)” Early AD” analysis, B) “Late AD” analysis and C) “Standard AD” analysis. All brain regions in this study are affected in AD, specifically by atrophy and neuronal loss, while only three brain regions in this study (EC, TC and FC) are affected by the additional accumulation of hallmark AD pathology (A_β_ and NFT). Genes perturbed across all brain regions may be markers of cell dysfunction in AD, while genes consistently perturbed in the EC, TC and FC but not in the CB may be associated with AD pathology. **MOSPD3** gene is the only gene DE in all brain regions of the “Early AD” analysis. No gene is DE across all brain regions in the “Late AD” analysis. Three (**ALDH2, FBLN2**, **METTL7A**) and nine genes (**FLCN, ASPHD1, ARL5A, GPR162, HBA2, PCID2, NDRG2, BEND3, RAP1Gap**) are consistently DE across all brain regions affected by hallmark AD pathology in the “Early AD” and “Late AD” analyses respectively.

### Differentially expressed genes in brain regions with hallmark AD pathology

The EC, TL and FC are all affected by hallmark AD pathology (amyloid and NFT’s), while the CB is known to be partially spared. Gene’s DE in the EC, TC and FC brain regions and not the CB, may identify hallmark AD pathology specific genes. Three (**ALDH2, FBLN2** and **METTL7A**) and nine (**FLCN, ASPHD1, ARL5A, GPR162, HBA2, PCID2, NDRG2, BEND3, RAP1Gap**) genes were significantly differentially expressed across the EC, TC and FC brain regions and not the CB brain region in the “Early AD” and “Late AD” analysis respectively.

### Gene Set Enrichment Analysis of differentially expressed genes

To understand the functional implications of DEG’s, GSEA was performed using the significant DEG list from all three analyses (“Early AD”, “Late AD” and “Standard AD”) and across all four brain regions, resulting in 12 enrichment result tables (provided in Supplementary Table 4). No biological pathway is significantly enriched across all four brain regions in the “Early AD”, “Late AD” or “Standard AD” analysis. However, when excluding the brain region often referred to spared by hallmark AD pathology (CB), the “**glutamate glutamine metabolism”** and “**gluconeogenesis and glycolysis**” pathways are the only pathways significantly enriched in the “Early AD” and “Late AD” analysis respectively. For the “Standard AD” analysis, excluding the CB brain region additionally identified “**mRNA processing**”, “**synaptic vesicle pathway**” and “**TNF-alpha**” pathways as significantly enriched in the remaining three brain regions.

### Summary of Weighted Co-Expression Network Analysis

Weighted gene co-expression analysis was performed on the FC and EC brain regions. We focused on these two brain regions as differential expression analysis identified an increased number of significant DEG’s in the FC brain region prior to AD symptoms and the EC is widely regarded as one of the first areas of the brain to be affected in AD. Network preservation and cross-tabulation statistics were calculated to identify co-expression networks that may be preserved or disrupted between the Control, AsymAD and AD subjects. Figure 5 illustrates the WGCNA module assignments and module preservation statistics, and Figure 6 shows the cross-tabulation statistics across phenotypes.

**Figure 5:**
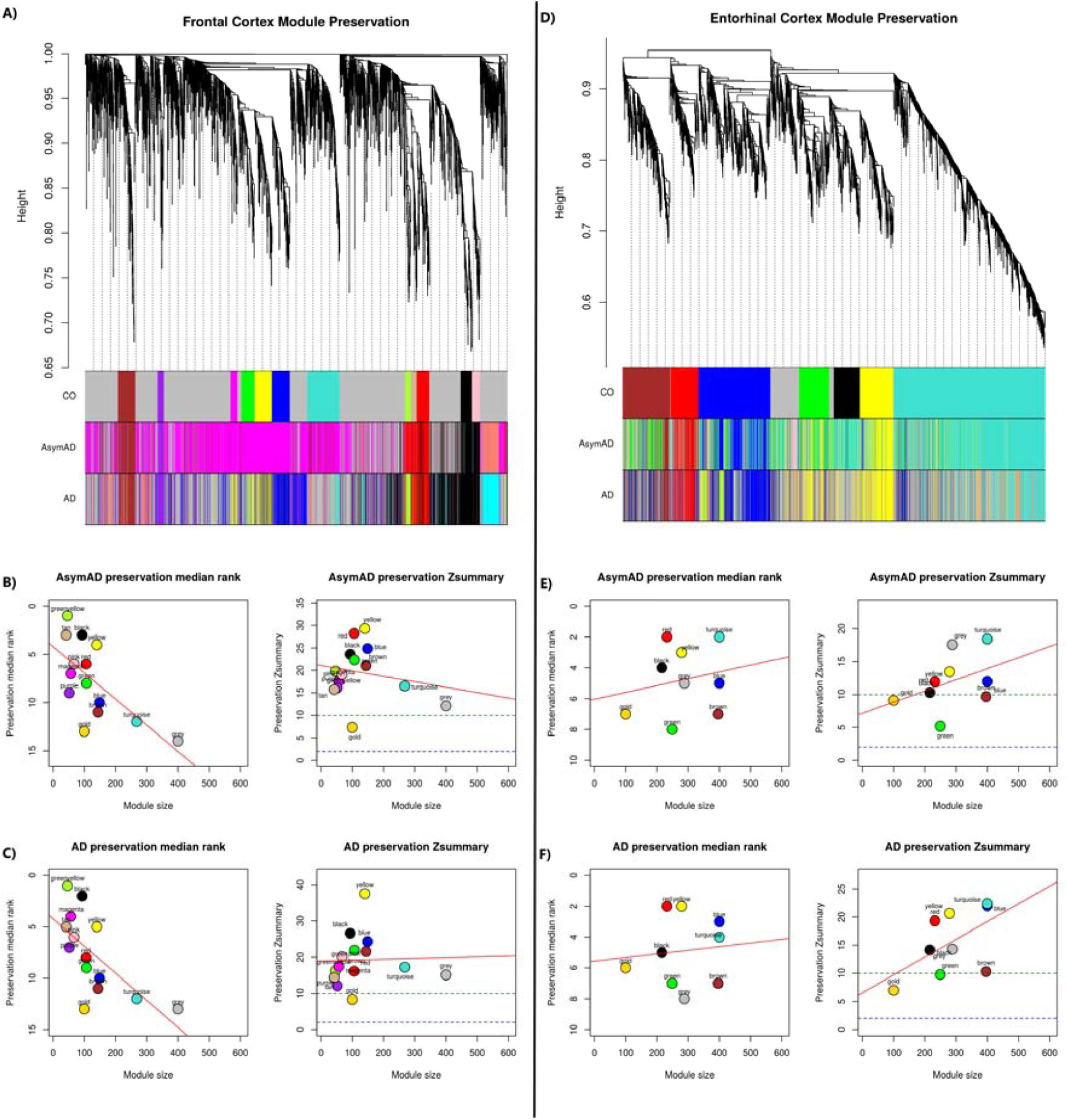
Hierarchical clustering of genes and module preservations statistics for the frontal cortex is illustrated in A-C) and entorhinal cortex in D-F). In brief, a co-expression network based on “signed” adjacency was independently created for all three phenotypes (control, AsymAD and AD group), topological overlap calculated, and hierarchical clustering used to group genes into modules. For the Hierarchical clustering plots, the y-axis represents the network distance with values closer to 0 indicating greater similarity of probe expression across the control group. The x-axis represents the modules in the control, AsymAD and AD group. The AsymAD and AD module colours are mapped to the control group, with the AsymAD and AD colour panel representing how well the control modules are preserved through the disease. The red line in the module preservation statistics (B, C, E, F) represents the correlation between module size and preservation statistics. The gold module represents 100 random genes, and the grey module represents uncharacterised genes. The FC preservation plots (B and C) suggest all modules in the control group are relatively preserved in the AsymAD and AD group. In contrast, the EC preservation plots (E and F) suggest the green module is not well preserved in the AsymAD and AD group and requires further investigation.

**Figure 6:**
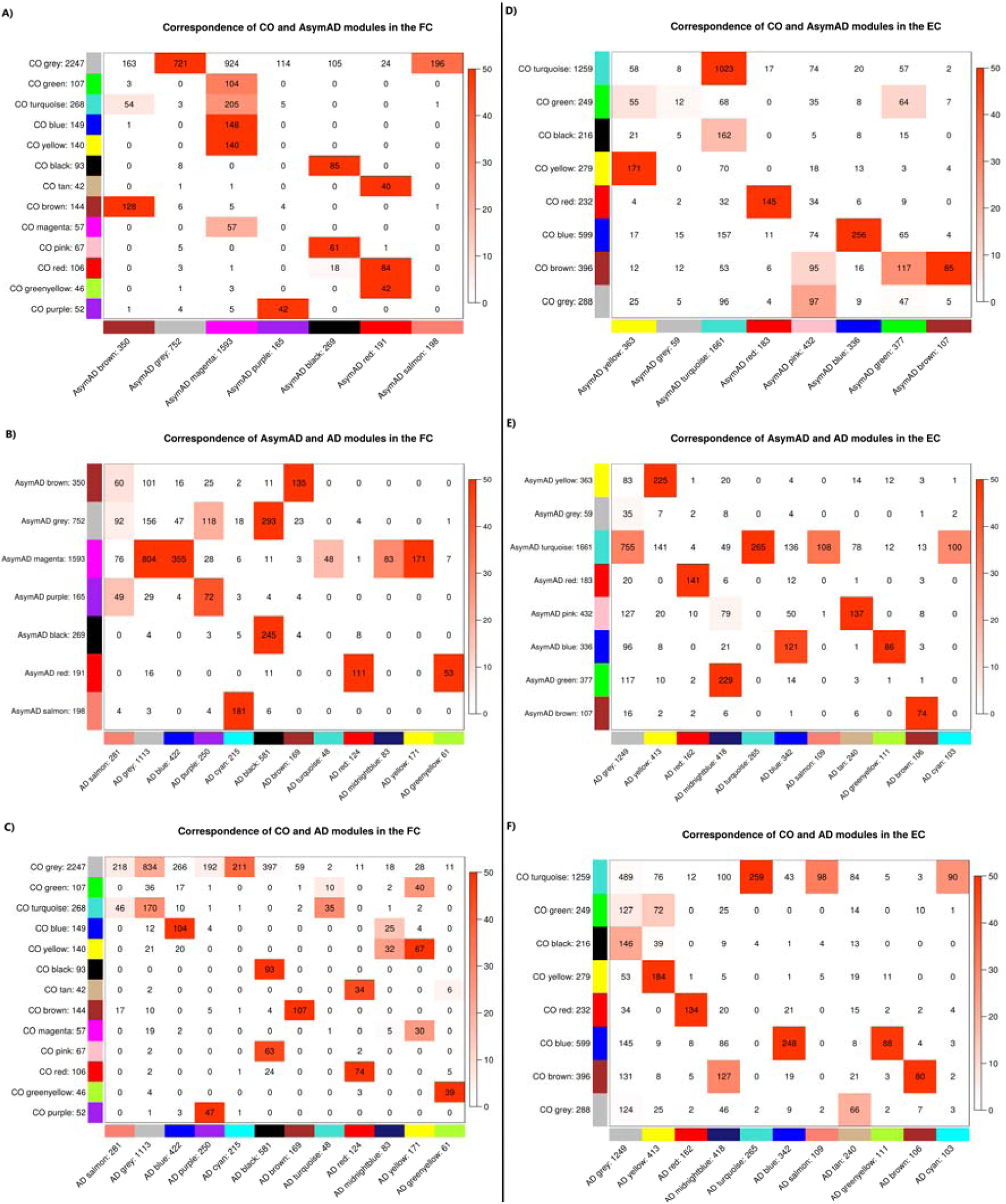
Illustrates the “module correspondence” between A) FC control and AsymAD group, B) FC AsymAD and AD group, C) FC control and AD group, D) EC control and AsymAD, E) EC AsymAD and AD group, and F) EC control and AD groups. The modules represent clusters of highly correlated genes which were calculated independently in each brain region and diagnosis group. The module colours in the AsymAD and AD group were assigned based on the gene overlap of the control module. The total number of genes within each module is indicated next to the module colour. The numbers in each cell represent the overlap of genes between modules, with increased red intensity cells indicating increased significant overlap based on Fisher’s exact test. This “module correspondence” plot provides a visual overview of how modules of highly correlated genes are preserved or disrupted between, control, AsymAD and AD groups. Module preservation statistics suggested the green module in the EC control group is not well preserved in the AsymAD and AD groups, indicating possible disruption to the co-expression network in this module. This “module correspondence” plot identifies the disrupted genes in the control green module synchronises with the genes of the AsymAD yellow module, identifying the yellow module for further investigation.

Co-expression analysis in the FC brain region identified 13, 7, and 12 modules within the control, AsymAD and AD groups respectively, while analysis in the EC identified 8, 8 and 11 modules within the control, AsymAD and AD groups respectively. GSEA analysis was performed for all fifty-nine modules to identify potential biological pathways the co-expressed genes may be involved with. A summary of the GSEA results on the co-expression module in the FC and EC is provided in Table 2 and Table 3 respectively, with complete GSEA results for the Control, AsymAD and AD groups in the FC and EC brain regions provided in Supplementary Tables 5, 6, 7, 8, 9, 10 respectively.

**Table 2:**
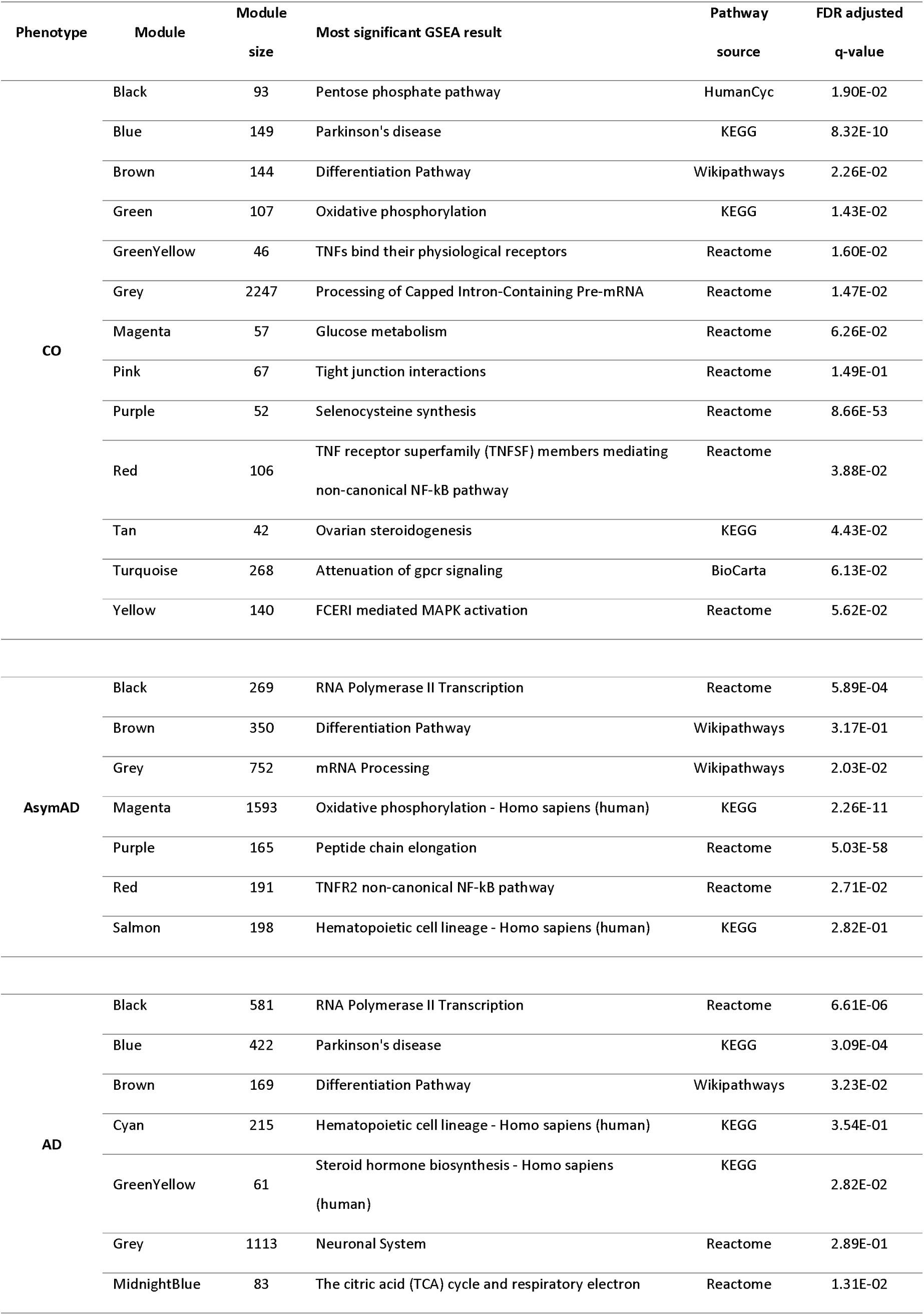
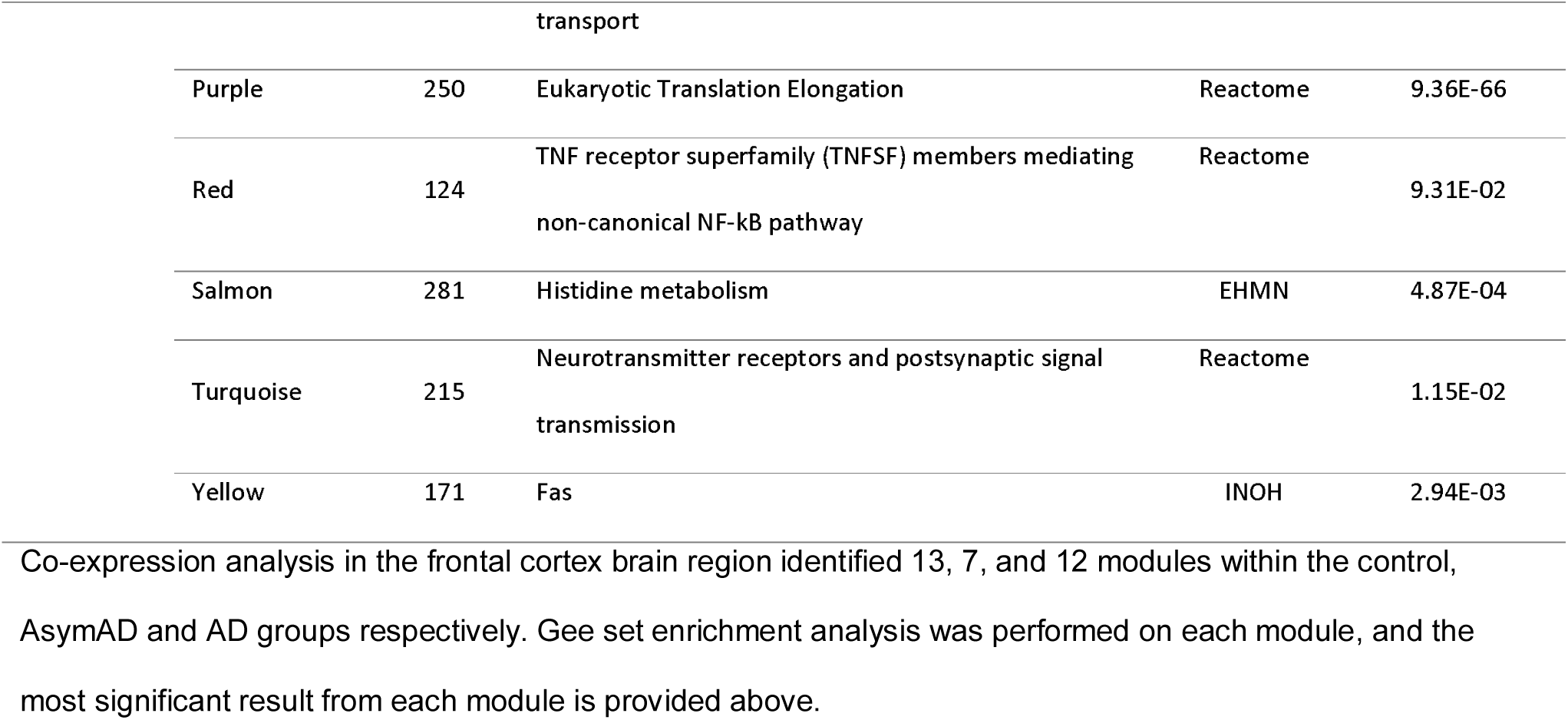
Summary of frontal cortex co-expression module GSEA results

**Table 3:**
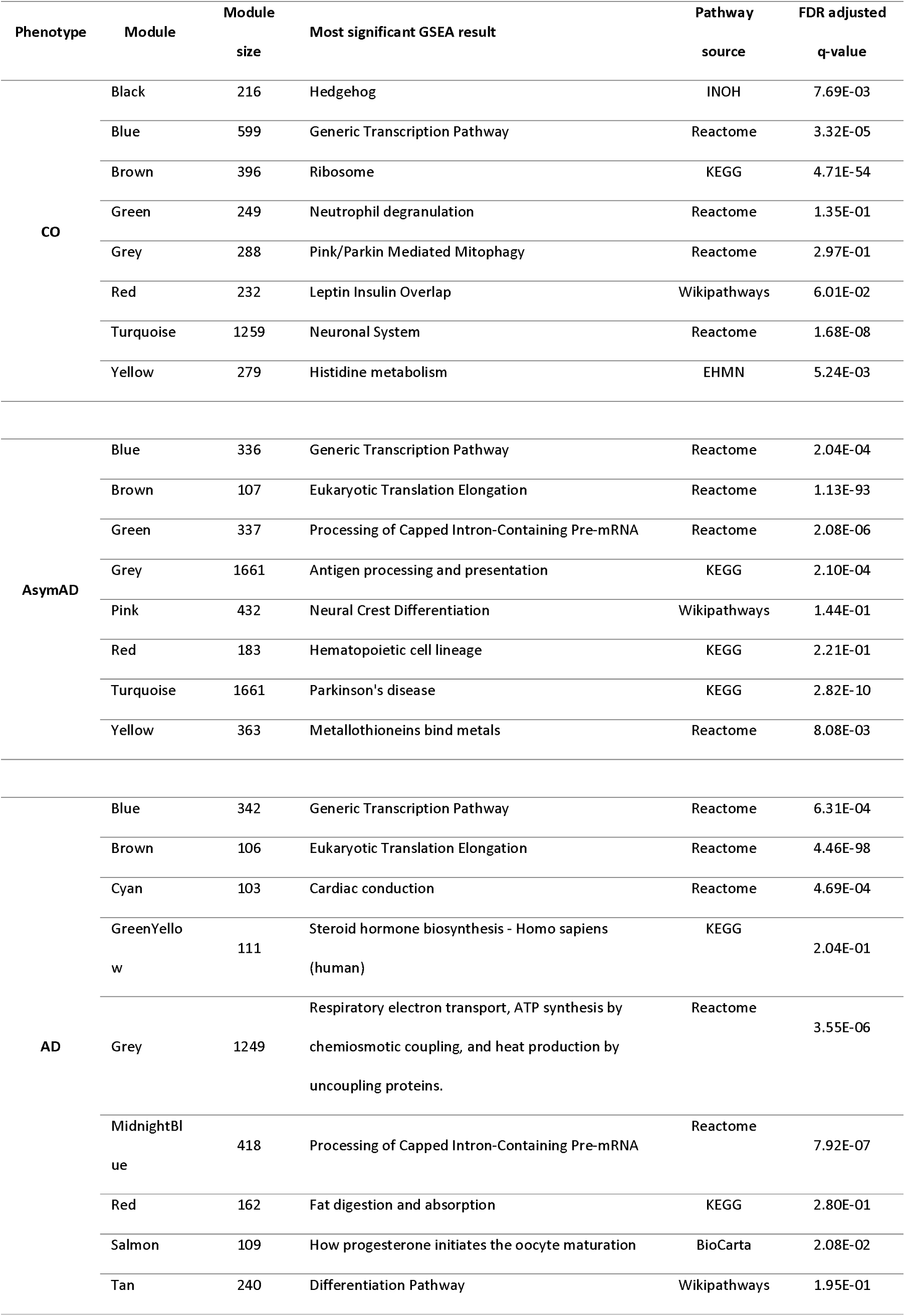
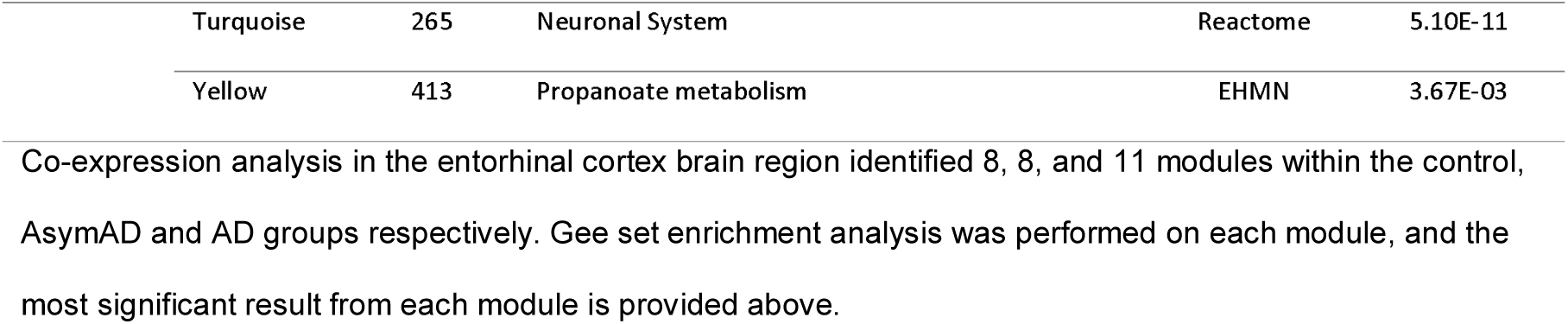
Summary of entorhinal cortex module co-expression results

| Phenotype | Module | Module size | Most significant GSEA result | Pathway source | FDR adjusted q-value |
| --- | --- | --- | --- | --- | --- |
| CO | Black | 216 | Hedgehog | INOH | 7.69E-03 |
|  | Blue | 599 | Generic Transcription Pathway | Reactome | 3.32E-05 |
|  | Brown | 396 | Ribosome | KEGG | 4.71E-54 |
|  | Green | 249 | Neutrophil degranulation | Reactome | 1.35E-01 |
|  | Grey | 288 | Pink/Parkin Mediated Mitophagy | Reactome | 2.97E-01 |
|  | Red | 232 | Leptin Insulin Overlap | Wikipathways | 6.01E-02 |
|  | Turquoise | 1259 | Neuronal System | Reactome | 1.68E-08 |
|  | Yellow | 279 | Histidine metabolism | EHMN | 5.24E-03 |
| AsymAD | Blue | 336 | Generic Transcription Pathway | Reactome | 2.04E-04 |
|  | Brown | 107 | Eukaryotic Translation Elongation | Reactome | 1.13E-93 |
|  | Green | 337 | Processing of Capped Intron-Containing Pre-mRNA | Reactome | 2.08E-06 |
|  | Grey | 1661 | Antigen processing and presentation | KEGG | 2.10E-04 |
|  | Pink | 432 | Neural Crest Differentiation | Wikipathways | 1.44E-01 |
|  | Red | 183 | Hematopoietic cell lineage | KEGG | 2.21E-01 |
|  | Turquoise | 1661 | Parkinson's disease | KEGG | 2.82E-10 |
|  | Yellow | 363 | Metallothioneins bind metals | Reactome | 8.08E-03 |
| AD | Blue | 342 | Generic Transcription Pathway | Reactome | 6.31E-04 |
|  | Brown | 106 | Eukaryotic Translation Elongation | Reactome | 4.46E-98 |
|  | Cyan | 103 | Cardiac conduction | Reactome | 4.69E-04 |
|  | GreenYellow | 111 | Steroid hormone biosynthesis - Homo sapiens (human) | KEGG | 2.04E-01 |
|  | Grey | 1249 | Respiratory electron transport, ATP synthesis by chemiosmotic coupling, and heat production by uncoupling proteins. | Reactome | 3.55E-06 |
|  | MidnightBlue | 418 | Processing of Capped Intron-Containing Pre-mRNA | Reactome | 7.92E-07 |
|  | Red | 162 | Fat digestion and absorption | KEGG | 2.80E-01 |
|  | Salmon | 109 | How progesterone initiates the oocyte maturation | BioCarta | 2.08E-02 |
|  | Tan | 240 | Differentiation Pathway | Wikipathways | 1.95E-01 |

|  |  |  |  |  |
| --- | --- | --- | --- | --- |
| Turquoise | 265 | Neuronal System | Reactome | 5.10E-11 |
| Yellow | 413 | Propanoate metabolism | EHMN | 3.67E-03 |

Co-expression analysis in the entorhinal cortex brain region identified 8, 8, and 11 modules within the control, AsymAD and AD groups respectively. Gse set enrichment analysis was performed on each module, and the most significant result from each module is provided above.

### Co-expression modules are weakly preserved in AsymAD and AD entorhinal cortex

Module preservation statistics were calculated for each brain region to identify co-expression networks that are weakly preserved through the course of the disease. Modules below a “preservation Zsummary” statistic of 10 and “preservation median rank” higher than the gold module (random 100 genes) are suggested to be weakly preserved. Module colours for the AsymAD and AD groups were mapped to the control module colours, allowing for changes and preservation in the co-expression networks to be observed as the disease progresses. The module colours assigned in the EC brain region are independently assigned to modules colours assigned in the FC brain region and therefore similar module colours across these two tissues bare no relation.

The FC “preservation Zsummary” statistics (Figure 5b and Figure 5c) suggests all modules from the control group are relatively well-preserved in the AsymAD and AD groups. In contrast, the EC “preservation median rank” statistics suggest the green control module is weakly preserved in AsymAD group (Figure 5e), and both the green and brown control modules are weakly preserved in the AD group (Figure 5f). In addition, the cross-tabulation statistics are also indicative of disruption to the EC green control module (Figure 6d). GSEA reveals the EC brown module in control, AsymAD and AD group is most significantly enriched for “**selenocysteine synthesis**” (control q-value=4.71e54, AsymAD q-value=5.89e-90, AD= 1.35E-96), suggesting this process is not significantly disrupted in AsymAD or AD subjects. In contrast, the EC control green module is significantly enriched (before multiple corrections) for **“neutrophil degranulation”** (p-value = 0.5e-4), **“TYROBP casual network”** (p-value = 2.5e-3) and the **“innate immune system”** (p-value = 2.7e-3), none of which are present in the green module of the AsymAD, suggesting these pathways may be disrupted in AsymAD subjects.

Clusters of co-expressed genes in both the FC and EC brain regions were enriched for specific cell types including neurons, astrocytes, oligodendrocytes and microglia (results not shown); however, we did not detect a disturbance in any cell type in AsymAD subjects.

### Frontal Cortex Co-expression network re-wired in AsymAD

Co-expression analysis identified 13 and 12 co-expressed modules in the control and AD subjects respectively. However, the AsymAD group exhibits 7 larger modules of highly co-expressed genes, suggesting the co-expression network is re-wired in the FC brain region in this intermediate stage of AD. The module preservation analysis suggested all modules within the control group are relatively preserved through the course of the disease, however, through cross-tabulation of the modules we observe subtle changes leading to a much larger magenta module in the AsymAD group. The biological processes associated with the magenta module changes from being enriched for “**glucose metabolism**” (q-value 6.26e-02) in the control group to “**oxidative phosphorylation**” (q-value = 2.26e-11), **Parkinson’s disease** (q-value = 5.12e-9), **electron transport chain** (q-value = 5.83e-9) and **Alzheimer’s disease** (q-value = 8.21e-9) in the AsymAD group. Then the large magenta module in the AsymAD group, branches into four new AD modules (blue, turquoise, midnightblue, and yellow), which are most enriched for **Parkinson’s disease** (q-value = 3.09e-4), **neurotransmitter receptors and postsynaptic signal transmission** (q-value = 0.01), **the citric acid (TCA) cycle and respiratory electron transport** (q-value = 0.01), and **fas signalling** (q-value=0.0030) respectively.

### Entorhinal Cortex yellow module enriched for all “Early AD” analysis DEG’s

The yellow module contained all genes identified as significantly DE in the “Early AD” analysis (**ALDH2**, **FBLN1** and **METTL7A**) and contained a large number of genes disrupted from the green module, which was the least preserved module through disease progression. Overall, this made the yellow module a prime candidate for further investigation. Gene set enrichment analysis of the yellow module in the AsymAD group reveals enrichment in “**fatty acid degradation**“ (q-value=0.03), “**glycerophospholipid metabolism**” (q-value=0.008), “**urea cycle and metabolism of arginine, proline, glutamate, aspartate and asparagine**” (q-value=0.05), “**astrocytic glutamate-glutamine uptake and metabolism**” (q-value=0.05) and “**neurotransmitter uptake and metabolism in glial cells**” (q-value=0.05), all of which were not previously enriched in the matched yellow module in the control group.

Protein-protein interaction analysis in the yellow control module generated six networks, with the largest containing 28 nodes and 30 edges, and identified **EGFR** gene as the only significant key hub gene (p-value=0.01). **APOE** was not a member of this network. Further PPI analysis in the AsymAD yellow module generated five networks, with the largest containing 71 nodes and 81 edges, and **EGRF** was still the key hub gene (p-value=0.007). In the equivalent AD yellow module, PPI analysis identified a single network generated with 284 nodes and 420 edges. This network contained far higher numbers of genes and now integrated the **APOE** gene as part of the network with **UBC** as the key hub (p-value=4.12e63). This suggests protein interactions in this yellow module increases gradually through the course of the disease, with up-regulated **EGRF** interacting with more genes in the AsymAD group when compared to controls, followed by significant changes occurring in the AD group where up-regulated **UBC** gene takes more of a central role.

## Discussion

### Transcriptomic perturbations suggest AsymAD subjects could be an intermediate stage between control and MCI/AD

This study hypothesises the samples we have labelled as “AsymAD” subjects are an intermediate state between healthy ageing and MCI/AD. The assignment of these samples to the AsymAD group was based on the fact that these individuals had no reported clinical record of dementia prior to death as indicated in the MRC-LBB database; however, upon autopsy, these samples were found to have low levels of hallmark AD pathology, i.e. BRAAK Staging >= 2. Furthermore, an independent expression study identified the **TRIL** gene as being highly correlated with AD neuropathology, specifically tau pathology [39]. Our study shows that the **TRIL** gene expression gradually increases from the Control to AsymAD, and then further increases in AD subjects (Figure 2a), and this expression pattern is only observed in brain regions known to be affected by hallmark AD pathology (amyloid and NFT’s), i.e. the EC, TC and FC, and not in the CB brain region. This observation suggests the phenotype assignments (controls, AsymAD, AD) are a suitable representation of three points in AD progression (assuming the AsymAD subjects are all prodromal AD), and as suggested by the **TRIL** gene expression pattern across brain regions and the fact the CB has been consistently reported to be partially spared from hallmark AD pathology (amyloid and NFT’s), even those with severe AD pathology [40], genes whose expression pattern differs significantly in the CB from that consistently seen in the EC, TC and FC tissues may be associated with hallmark AD pathology.

### MOSPD3 gene is perturbed in the brains of AsymAD and blood of AD subjects

We identify **MOSPD3** as the only significant DE gene which is consistently down-regulated across all four brain regions in the AsymAD subjects suggesting this may be an early marker of cell dysfunction in AD. The **MOSPD3** gene encodes for a Motile Sperm Domain Containing 3 protein [provided by RefSeq, Jul 2008] and has been reported to be significantly down-regulated (p-value = 6.47E-05) in the blood of AD subjects when compared to MCI subjects [41]. This suggests MOSPD3 gene expression is significantly decreased in the brain before clinical signs of AD are apparent, however, blood gene expression levels are only significantly decreased after clinical signs of AD are apparent. It is difficult to interpret the biological relevance of this gene in AD, and further investigation is required.

### Genes perturbed in brain regions affected explicitly by hallmark AD pathology may be associated with plaques and tangles, providing new therapeutic targets

Many molecular and cellular changes occur in AD brains including nerve cell death, atrophy, loss of neurons and accumulation hallmark AD pathology, specifically plaques and tangles. However, not all brain regions are affected to the same degree. The CB, which only accounts for 10% of the brain but contains over 50% of the brains total neurons, is often regarded as being partially spared from AD as plaques and tangles are generally not reported [40] [42], and in this study are free from hallmark AD pathology in both AsymAD and AD subjects. For subjects with hallmark AD pathology (BRAAK >=2, AsymAD and AD), genes significantly and consistently perturbed across the EC, TC and FC tissues that are not or are significantly reversed in the CB, may be associated with hallmark AD pathology, although, it remains unclear if these genes are causative or a response to the pathology itself.

We identified a total of 15 genes (**ALDH2, FBLN2**, **METTL7A**, **FLCN**, **ASPHD1, ARL5A, GPR162, HBA2, PCID2, NDRG2, BEND3, RAP1Gap, GPM6B, ANKEF1** and **NPC2**) with expression patterns suggestive of association with hallmark AD pathology. Previous studies have already demonstrated an increased expression of **ALDH2** accelerated neurodegeneration and increased the accumulation of hyperphosphorylated tau protein [43] in mice, while another demonstrated **NDRG2** might play a role in generating Aβ [44]. Collectively, the 15 genes are not significantly enriched to be involved with any biological pathway; however, individually, these genes may play an essential role in the pathological aspect of AD and may provide new therapeutic targets for disease intervention.

### Individuals with milder disease (early BRAAK pathology) show increased changes in the frontal cortex compared to the entorhinal cortex

The molecular changes in AD may initially begin in the FC, a region involved in working memory, as there were relatively more changes in the FC of mild pathology AD cases (AsymAD) than the EC region. This mirrors changes described in a longitudinal study involving ageing controls, where positron emission tomography (PET) scans were used to detect increased activity in the medial frontal cortex and decrease activity in the temporal lobe brain region in subjects who subsequently acquired cognitive impairment [45]. In addition, a higher degree of atrophy has also been detected in the FC than the temporal lobe brain region in MCI when compared to AD [46]. Our observations provide further evidence to suggest that brain perturbations at the molecular/transcriptomic level may initially occur in the FC before the presentation of more severe clinical symptoms consistent with a diagnosis of probable AD.

At the later point of the disease when clinical signs of AD are present, we find that the most substantial number of transcriptomic changes occur in the EC, followed by the TC, FC and only minor changes in the CB. This observation matches the common route AD neuropathology is seen to spread through the brain. Furthermore, we detect more DEG in the “Late AD” analysis compared to “Early AD” analysis, signifying more genes are disrupted in the later stage of the disease when the clinical symptoms of cognitive impairment are apparent.

### Neutrophil, TYROBP network and the innate immune system disrupted in Asymptomatic AD

Co-expression analysis of the EC brain region identified a green module of highly co-expressed genes which is disrupted in the AsymAD and AD subjects according to both module preservation statistics and cross-tabulation analysis. This green module is significantly enriched for “**neutrophil degranulation**”, “**TYROBP casual network**” and the “**innate immune system**” processes in the control subjects, but not in the AsymAD or AD subjects, suggesting these pathways are most likely disrupted during the disease. Disturbance in TYROBP and Immune system pathways have been widely accepted in AD [47] [15], and a previous mouse study demonstrated disruptions in neutrophil levels impact memory loss and neurological features of AD [48]. We now suggest these pathways are specifically perturbed in the EC brain region early in the disease when hallmark AD pathology exists but clinical symptoms of AD are absent.

### Disruption in brain energy pathways is detectable early in the disease

Co-expression analysis of the FC identifies disruptions in the “**glucose metabolism**”, “**glucogenesis**” and“**oxidative phosphorylation**” processes in the AsymAD group, while DE analysis identified disruption in the “**gluconeogenesis and glycolysis**” pathway in the AD subjects. The brain critically relies on a constant supply of energy which is known to be generated by glycolysis followed by oxidative phosphorylation. Changes in the brain energy pathways have been widely accepted in AD [49] [50], with a general decrease in glycolysis suggested to be a result of decreased brain functionality. Here we demonstrate disruptions in the energy pathway are detectable early in the disease, in subjects with low levels of AD pathology.

### The Glutamate-Glutamine Cycle is disturbed in AsymAD and AD subjects

Gene set enrichment analysis on DEGs identified the “**glutamate-glutamine cycle**” as the only biological pathway significantly perturbed across all brain regions in the AsymAD subjects. Furthermore, co-expression analysis of the EC brain regions was indicative of disruptions to the “**urea cycle and metabolism of arginine, proline, glutamate, aspartate and asparagine**” and “**astrocytic glutamate-glutamine uptake and metabolism**” in AsymAD and AD subjects, further confirming a possible disruption in glutamate-related activities in the brain.

Astrocytes are the most common form of neuroglial cells in the brain, and its primary function is to protect neurons against excitotoxicity by converting excess ammonia and glutamate to glutamine through the glutamate-glutamine cycle. Glutamate is the principal excitatory neurotransmitter in the brain and plays a vital role in linking carbohydrate and amino acid metabolism via the tricarboxylic acid (TCA) cycle. Glutamate is also a precursor of γ-aminobutyric acid (GABA) which binds and inhibits neuron activity; hence, an accumulation of glutamate can cause failures in synaptic connectivity, leading to deficient cognition and memory [51]. A disruption in the glutamate-glutamine cycle would have a severe knock-on effect on many other biological pathways, including a disruption in amino acid metabolism which could explain the enrichment of “**urea cycle and metabolism of arginine, proline, glutamate, aspartate and asparagine**” in our results as well. In addition, glutamate stimulates astrocytes to derive energy from oxidative and glycolytic pathways, both of which have been identified as disrupted in AsymAD subjects.

The genes enriched in this pathway were all significantly up-regulated, indicating an overactive cycle. This could be part of the brain defence mechanism in preventing accumulation of brain glutamate levels or a broken cycle which is consistently being overactive, leading to decreased levels of brain glutamate, a phenomenon observed in AD subjects [52]. Targetting this pathway for AD treatment is extraordinarily complex and challenging as over inhibition or excitation may lead to increased levels of glutamate and glutamine respectively, both of which can be neurotoxic at high levels. Therapeutic compounds affecting the “**glutamate-glutamine cycle**” have already been identified, such as memantine, which is already a clinically established therapeutic drug used to for the symptomatic treatment of AD, which blocks N-methyl-d-aspartate (NMDA) receptors [53], essentially preventing excitotoxicity caused by neurotransmitters such as glutamate and ultimately increasing cognition temporarily.

The glutamate-glutamine cycle has been previously suggested to be disrupted in AD [54], along with many other central nervous system disorders including Huntington’s disease and Amyotrophic Lateral Sclerosis (ALS) [55]. Through this study, we now demonstrate this is one of the earliest biological pathways perturbed across all brain regions in AD, before clinical symptoms of AD are apparent, which can have a knock-on effect on other biological pathways also observed to be disrupted in the disease. Clinically established drugs to relieve AD symptoms already interact with this pathway and could also be effective in the asymptomatic period to prolong cognitive impairment, although clinical identification and measuring effectiveness in AsymAD subjects would be a challenge in itself.

### Co-expression network changes indicate a shift from “cell proliferation” in AsymAD subjects to “removal of amyloidogenic proteins” in AD subjects

Protein-protein interactions identified **EGFR** as a key hub gene in both the control and AsymAD groups; however, it achieves more connections with neighbouring proteins in the AsymAD group, suggesting a possible increase in the **EGFR** activity. The **EGFR** gene is up-regulated in the AsymAD group and encodes for a transmembrane glycoprotein that binds to epidermal growth factor, leading to cell proliferation. In contrast, **EGFR** is replaced by **UBC** as the key hub gene in AD subjects, indicating it may play a more central role in the disease once accumulation of hallmark AD pathology is at a level where clinical symptoms are apparent. The **UBC** gene is significantly up-regulated in the EC of AD subjects and is considered a stress gene which encodes for polyubiquitin precursor protein, a member of the ubiquitin-proteasome system (UPS) which removes toxic proteins and impacts on the amyloidogenic pathway of amyloid precursor protein (**APP**) processing that generates Abeta [56]. A previous AD study had also observed **UBC** as a novel key hub gene and demonstrated **UBC** knockout models in C. elegans accelerated age-related AB toxicity [57]. Effectively, a portion of the co-expression network may have a central role involved in cell proliferation in control subjects, with increased activity in AsymAD subjects, followed by a shift towards the removal of toxic proteins such as amyloid beta in AD subjects.

### Limitations

We cannot exclude the fact AsymAD group may represent a heterogeneous group consisting of cognitively normal, MCI, mixed dementia and AD subjects. It remains unclear these AsymAD subjects would remain free from clinical symptoms of dementia with longer survival and can be argued to be a possible extension to general ageing. However, the extent of BRAAK staging in AsymAD subjects was at a level consistently found with early cognitive impairment, and therefore, we make the strong assumption that these subjects are more likely to be prodromal AD rather than an extension of natural ageing. As AsymAD subjects are extremely rare, hence the low sample numbers in this study, larger AsymAD cohorts are required for better discovery and to validate our findings.

### Conclusion

We believe this is the first study to explore the emergence of transcriptomic changes in the human brain from normal ageing through to mild AD pathology and diagnosis of AD. Using DE analysis, coupled with a “systems-biology” approach, we were able to detect disturbances in the energy pathways and the “**glutamate-glutamine cycle**” in the brains of subjects with mild and severe AD pathology. We found that changes in the FC brain region dominate in mild pathology, but are greater in the EC in subjects with more severe pathology, thus mirroring the changes in aggregate spread in AD. This study provides new insight into the earliest biological changes occurring in the brain prior to AD diagnosis while providing new potential therapeutic targets.

## Acknowledgements

This study presents independent research supported by the NIHR BioResource Centre Maudsley at South London and Maudsley NHS Foundation Trust (SLaM) & Institute of Psychiatry, Psychology and Neuroscience (IoPPN), King’s College London. The views expressed are those of the author(s) and not necessarily those of the NHS, NIHR, Department of Health or King’s College London. We gratefully acknowledge capital equipment funding from the Maudsley Charity (Grant Ref. 980) and Guy’s and St Thomas’s Charity (Grant Ref. STR130505. Tissue samples were supplied by the London Neurodegenerative Diseases Brain Bank, which receives funding from the MRC and as part of the Brains for Dementia Research programme, jointly funded by Alzheimer’s Research UK and Alzheimer’s Society.

RJBD and SJN are supported by 1. Health Data Research UK, which is funded by the UK Medical Research Council, Engineering and Physical Sciences Research Council, Economic and Social Research Council, Department of Health and Social Care (England), Chief Scientist Office of the Scottish Government Health and Social Care Directorates, Health and Social Care Research and Development Division (Welsh Government), Public Health Agency (Northern Ireland), British Heart Foundation and Wellcome Trust. 2. The National Institute for Health Research University College London Hospitals Biomedical Research Centre.

